# Development of amplicon deep sequencing markers and data analysis pipeline for genotyping multi-clonal malaria infections

**DOI:** 10.1101/121426

**Authors:** Anita Lerch, Cristian Koepfli, Natalie Hofmann, Camilla Messerli, Stephen Wilcox, Johanna H. Kattenberg, Inoni Betuela, Liam O’Connor, Ivo Mueller, Ingrid Felger

**Author notes:** current affiliations: JHK: Institute of Tropical Medicine, Antwerp, Belgium; IM: Institut Pasteur, Paris, France.

## Abstract

Amplicon deep sequencing permits sensitive detection of minority clones and improves discriminatory power for genotyping multi-clone *Plasmodium falciparum* infections. Such high resolution is needed for molecular monitoring of drug efficacy trials. Targeted sequencing of molecular marker *csp* and novel marker *cpmp* was conducted in duplicate on mixtures of parasite culture strains and 37 field samples. A protocol to multiplex up to 384 samples in a single sequencing run was applied. Software “HaplotypR” was developed for data analysis. *Cpmp* was highly diverse (H_e_=0.96) in contrast to *csp* (H_e_=0.57). Minority clones were robustly detected if their frequency was >1%. False haplotype calls owing to sequencing errors were observed below that threshold. To reliably detect haplotypes at very low frequencies, experiments are best performed in duplicate and should aim for coverage of >10’000 reads/amplicon. When compared to length polymorphic marker *msp2*, highly multiplexed amplicon sequencing displayed greater sensitivity in detecting minority clones.

## Introduction

In infection biology of malaria as well as of many other pathogens, detection of minority clones is a crucial task. In areas of high malaria transmission, most infected hosts harbour multiple clones of the same *Plasmodium* species. To better understand the epidemiology and infection dynamics of malaria, individual parasite clones are tracked over time to measure the acquisition, elimination and persistence of individual clones in a human host. The incidence of new clones per host serves as surrogate measure for the exposure of an individual and for the transmission intensity in a population (1). Identification of new infections is also crucial in clinical trials of antimalarial drugs, where persisting clones need to be distinguished from new clones in post-treatment samples from patients with recurrent parasitaemia (2,3). For such diverse applications, genotyping methods based on length polymorphic markers had been applied for decades, particularly by targeting microsatellite markers or genes encoding parasite surface antigens such as merozoite surface proteins 1 and 2 (*msp1, msp2*)(4,5).

Despite their wide use in many malaria research laboratories, length polymorphic markers have important limitations. For example, microsatellite typing suffers from frequent occurrence of stutter peaks, possibly resulting from polymerase slippage on stretches of simple tandem repeats. A cut-off requirement for a minimal peak height (e.g. 33% of the predominant peak (6)) is required to prevent scoring of artefact fragments. However, this cut-off makes it impossible to detect minority clones falling below the selected threshold. Another limitation of length polymorphic marker, particularly the highly polymorphic parasite surface antigens, consists in the usually large size differences between alleles. Major size differences lead to bias in amplification, preferring the shorter PCR fragments in samples that concurrently contain multiple *P. falciparum* infections (7).

Deep sequencing of short amplicons has the potential to overcome some of the shortfalls of length polymorphic genotyping markers, in particular the influence of fragment length of a marker on the detectability of minority clones. Earlier studies used two different approaches for genotyping of *P. falciparum* and *P. vivax* by amplicon deep sequencing: (i) Sequencing of the classical length polymorphic genotyping markers, such as *P. falciparum msp1* and *msp2* (8). Alternatively, sequencing targeted non-repetitive regions that harbour extensive single nucleotide polymorphism (SNP), such as the *P. falciparum* circumsporozoite protein (*csp*) or *P. vivax msp1* (9,10). The strength of these approaches is that all SNPs within an amplicon are linked by a single sequence read, leading directly to haplotype identification. (ii) Sequencing of multiple loci of genome-wide distribution, whereby each locus comprises one SNP (11). This latter approach is particularly suited for population genetic investigations, as these loci are not linked. The downside is that the haplotype of each infecting clone has to be reconstructed, which is difficult or even impossible for samples with a high number of co-infecting clones per host (12). Thus, genotyping of samples containing multi-clone infections remains an unresolved challenge when multiple genome-wide loci are targeted.

In previous studies, amplicon deep sequencing was performed on two platforms, 454/Roche or Ion Torrent. In the past these technologies have produced longer sequences than the 37bp reads obtained by the Illumina sequencing platform. Now Illumina MiSeq generates reads of up to 600bp length (Illumina, MiSeq Reagent Kit v3). Sequencing error rates of 454/Roche and Ion Torrent technologies were high, owing to insertion and deletion (indel) errors occurring predominantly in homopolymeric regions (13–15). Illumina sequencing is less susceptible to indel errors and has an overall smaller error rate (15).

The present report outlines a strategy and protocols for identifying highly diverse markers for SNP-based genotyping of *P. falciparum* by amplicon sequencing. The primary aim was to thoroughly assess the analytical sensitivity and specificity of amplicon sequencing in detecting minority clones. In epidemiological studies involving hundreds of samples sequencing costs per sample are crucial. Therefore we designed a highly multiplexed protocol, allowing sequencing of up to 384 barcoded *P. falciparum* amplicons in a single Illumina MiSeq run. Because multiple concurrent *P. falciparum* clones may differ greatly in density, sequencing analysis strategies need to identify alleles of very low abundance. To distinguish true minority clones from sequencing errors, quality checks were designed based on replicates of samples and integrated into the sequence analysis pipeline. The newly created data analysis software package was validated using experimental mixtures of *P. falciparum in vitro* culture strains, and tested on field samples.

## Results

### Marker selection

A protocol for deep sequencing and data analysis was developed for two molecular markers, namely the *P. falciparum csp* gene (PF3D7_0304600) and gene PF3D7_0104100, annotated in the malaria sequence data base PlasmoDB as “conserved *Plasmodium* membrane protein” (*cpmp*). Results from these two markers were compared with classical length polymorphic genotyping using the highly diverse marker *msp2*. Sizes of *msp2* fragments amplified for genotyping range from 180 to 515 bp in PNG using published primers (Table S1). Marker *csp* has been used for deep sequencing in the past (9) and the previously published primers (Table S1) were used. The *csp* amplicon spans the T-cell epitope of the circumsporozoite protein from nucleotide position 858 to 1186 of the 3D7 reference sequence.

The newly validated marker *cpmp* was identified by calculating heterozygosity in 200 bp windows of 3’411 genomic *P. falciparum* sequences from 23 countries available from the MalariaGEN dataset (16). Genes from multi-gene families or regions of poor sequence alignments, often caused by length polymorphism of intragenic tandem repeats, were excluded from the list of potential markers. A 430bp fragment of *cpmp* spanning nucleotide positions 1895 to 2324 scored highest in expected heterozygocity (H_e_) and was prioritized as candidate for a highly diverse amplicon sequencing marker. He in the worldwide dataset was 0.93 for *cpmp* compared to 0.86 for *csp* (Table 1, Figures S1 and S2). Genomes originating from Papua New Guinea (PNG) revealed 9 haplotypes in 22 genomes for *cpmp* and 3 haplotypes in 30 genomes for *csp*.

**Table 1:** Diversity of markers *cpmp* and *csp* based on 3’411 genomes of the MalariaGen dataset.

| Marker | $H_e$<sup>1</sup> | No. of SNPs | Fragment size<sup>2</sup> | No. of Haplotypes |
| --- | --- | --- | --- | --- |
| *cpmp* | 0.930 | 20 | 383 | 82 of 980<sup>3</sup> |
| *csp* | 0.857 | 40 | 287 | 77 of 1323<sup>3</sup> |

<sup>1</sup> Expected Heterozygosity.
<sup>2</sup> Fragment size without primer sequence.
<sup>3</sup> From 3411 genomes only genomes with non-ambiguous SNP calls in selected region were used.

### Assessment of Sequence Quality

*Csp* and *cpmp* amplicons from 37 field samples and 13 mixtures of *P. falciparum* culture strains HB3 and 3D7 were sequenced on Illumina MiSeq in paired-end mode. A total of 5’810’566 paired raw sequences were retrieved. Of all reads, 326’302 mapped to the *phiX* reference sequence. 4’989’271 paired sequence reads were successfully de-multiplexed to yield a set of amplicon sequences per individual sample. 4’411’214 reads could be assigned to individual amplicons. Median sequence coverage over all sequenced samples was 1’490 for *cpmp* (1^st^ and 3^rd^ quartiles: [537, 2183]) and 731 for *csp* (1^st^ and 3^rd^ quartiles: [524, 1092]). The slight discrepancy was a result of the pooling strategy used (Supplementary File S1).

The quality of the sequence run was assessed by investigating the sequencing error rate in sequence reads of the spiked-in *phiX* control. The mean mismatch rate per nucleotide of *phiX* control reads with respect to the *phiX174* genome was 5.2% (median 0.34%). The mismatch rate increased towards the end of sequence reads, up to 11% for forward reads and 54% for reverse reads (Figure S3). To censor regions of high mismatch rates, forward and reverse sequence reads were trimmed before any further analyses to a length of 240 and 170 nucleotides, respectively. After trimming, the mean mismatch rate per nucleotide of *phiX* control reads was 0.50%.

As further quality check, the sequencing error rate was assessed in sequences of Linkers F and R (Figure S4). These linker sequences never get amplified but are joined to the product in PCR, therefore any mismatch detected in these stretches will derive from either sequencing or initial primer synthesis. The average number of sequence mismatches in this part was 0.12% per sample per nucleotide (Table S2, Figure S5). The sequencing error rate also was assessed in regions corresponding to the primers of each marker (Figure S4). Mismatches with respect to the known sequences of the PCR primers may derive from amplification errors or from errors in sequencing or primer synthesis during preparation of the sequencing library. The average number of mismatches in the primer regions was 0.28% for *cpmp* and 0.71% for *csp* per nucleotide per sample (Figure 1, Table S2).

**Figure 1:**
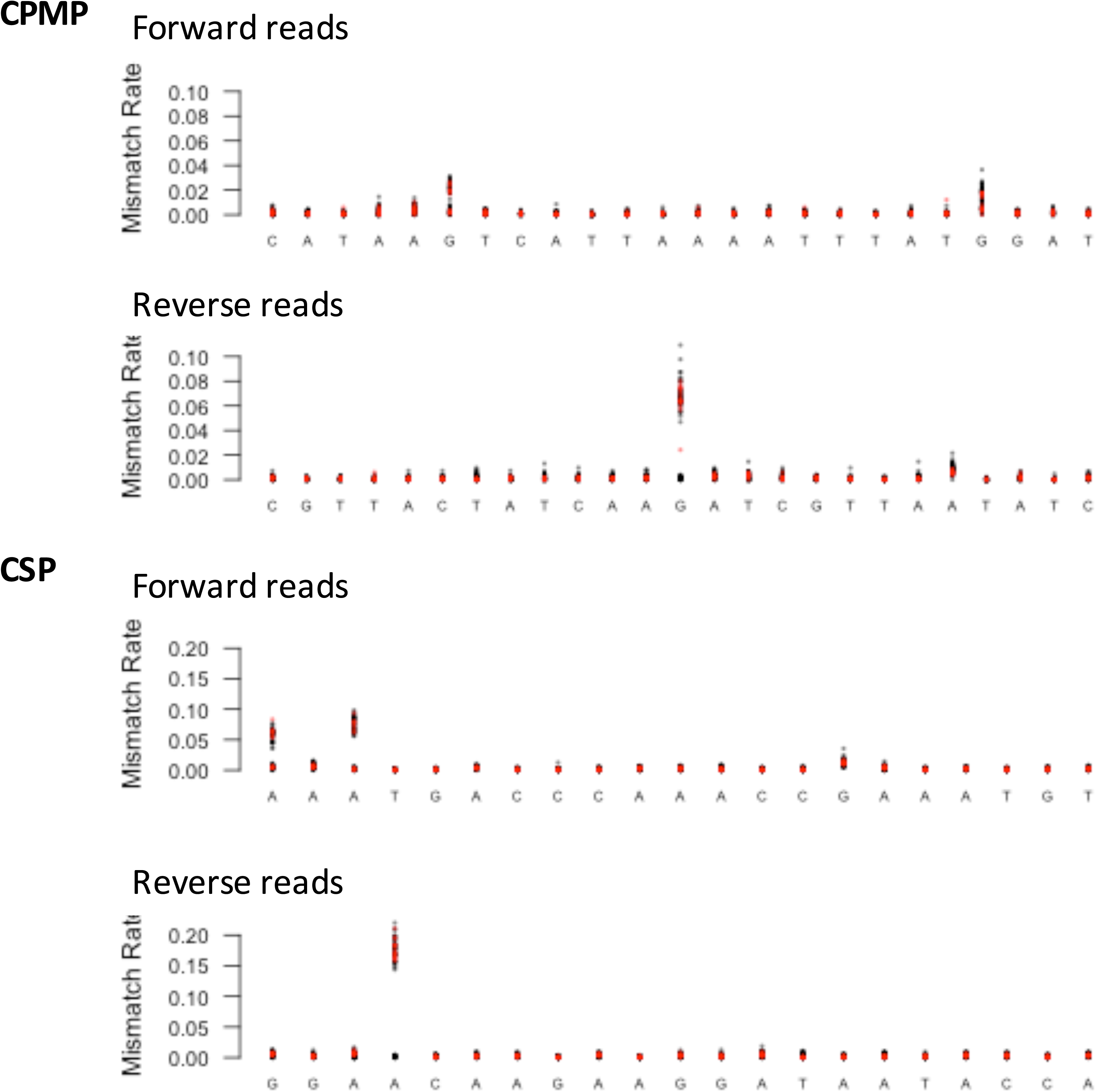
Mismatch rate per nucleotide position in forward and reverse primers of markers *cpmp* and *csp*. Data derived from all samples analysed. Each data point represents the mean observed mismatch rate of all reads from an individual sample. Red data points: control samples (*P. falciparum* culture strains); black data points: field samples; X-axis: nucleotides of forward and reverse primers (5’ to 3’); Y-axis: mismatch rate with respect to the known primer sequences.

Finally, the sequencing error rate was assessed in amplicons obtained from various mixtures of *P. falciparum* culture strains HB3 and 3D7. Potential sources of mismatches with respect to the reference sequence of strains 3D7 and HB3 include amplification error, sequencing error and errors due to de-multiplexing of samples (17). The average number of sequence mismatches after trimming to lengths of 240 and 170 nucleotides respectively for forward and reverse reads was 0.38% for *cpmp* and 0.46% for *csp* (Figure 2, Table S2). This equales to 1-2 mismatches per read of 310 nucleotides. On average 87.5% of reads for *cpmp* and 85.5% for *csp* from mixtures of strains HB3 and 3D7 contained ≤2 mismatches per read with respect to the strains’ reference sequences. Together the analyses of *phiX* and HB3/3D7 sequences indicated an intrinsic sequence error rate of 0.4-0.5%. The error rate of the linker sequence suggested that one third of these errors were sequencing errors, while two thirds were amplification errors.

**Figure 2:**
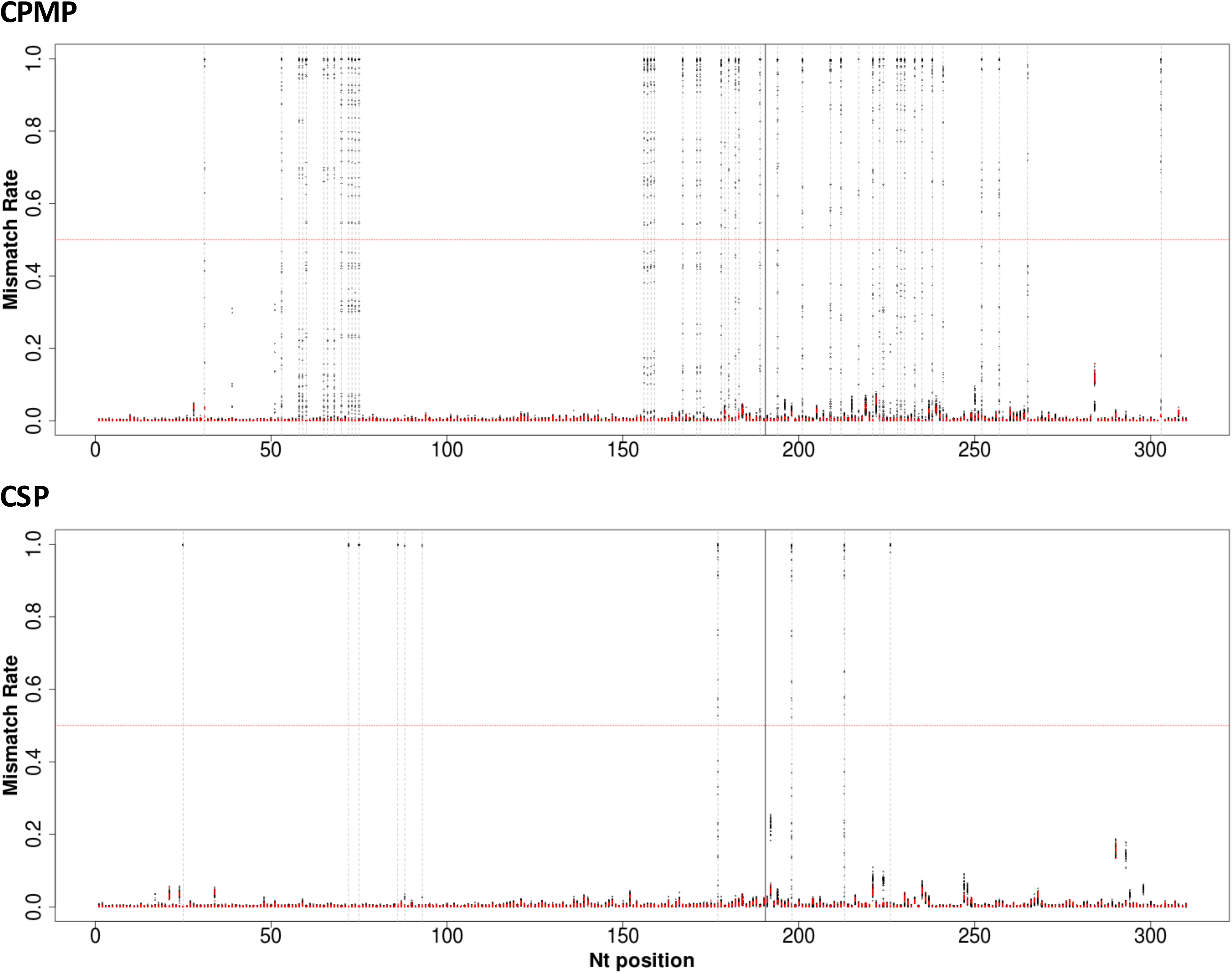
Mismatch rate per nucleotide position derived from all samples sequenced for markers *cpmp* and *csp*. Each data point represents the mean observed mismatch rate observed in all reads of one sample at the respective nucleotide position. Red data points: control samples (*P. falciparum* culture strains); black data points: field samples; X-axis: nucleotide position in sequenced fragment; Y-axis: mismatch rate with respect to the reference sequence (for control samples: sequences of strains 3D7 and HB3, for field samples, 3D7 sequence); dashed grey lines represent SNPs with a mismatch rate of >0.5 in >1 sample; red dotted horizontal line indicates a mismatch rate of 0.5; solid black vertical line: position of concatenation of forward and reverse reads.

### Limit of detection assessed in serial dilutions of parasite culture

To test the ability to genotype also blood samples of low parasite density, serial dilutions of *P. falciparum* strain 3D7 over 5 orders of magnitude (5-50’000 parasites/μl) were sequenced (Table S3). Using 1 μl of template DNA for PCR, dilutions harbouring 5 and 50 parasites/μl were detected, but with coverage below 550 reads. The pooling strategy applied (Supplementary File S1) did not fully counterbalance lower amounts of amplicon.

### Assessment of minority clone detectability

Defined mixtures of *P. falciparum* strains HB3 and 3D7 were sequenced to assess the detectability of minority clones under controlled conditions. The minority clone was detected in all tested dilution ratios up to 1:3000 (Table 2). Reads comprising obvious PCR artefacts (indels and chimeras) were detected in these mixtures up to a frequency of 0.48% for marker *cpmp* and 6.2% for *csp*. Up to 8.4% of reads for *cpmp* and 10.8% for *csp* were singletons or failed to cluster with 3D7 or HB3 haplotypes. This proportion of reads is therefore most likely an estimate of the cumulative background noise of the methodology. These reads fell below the default cut-off criteria (details below) and were thus excluded.

**Table 2:**
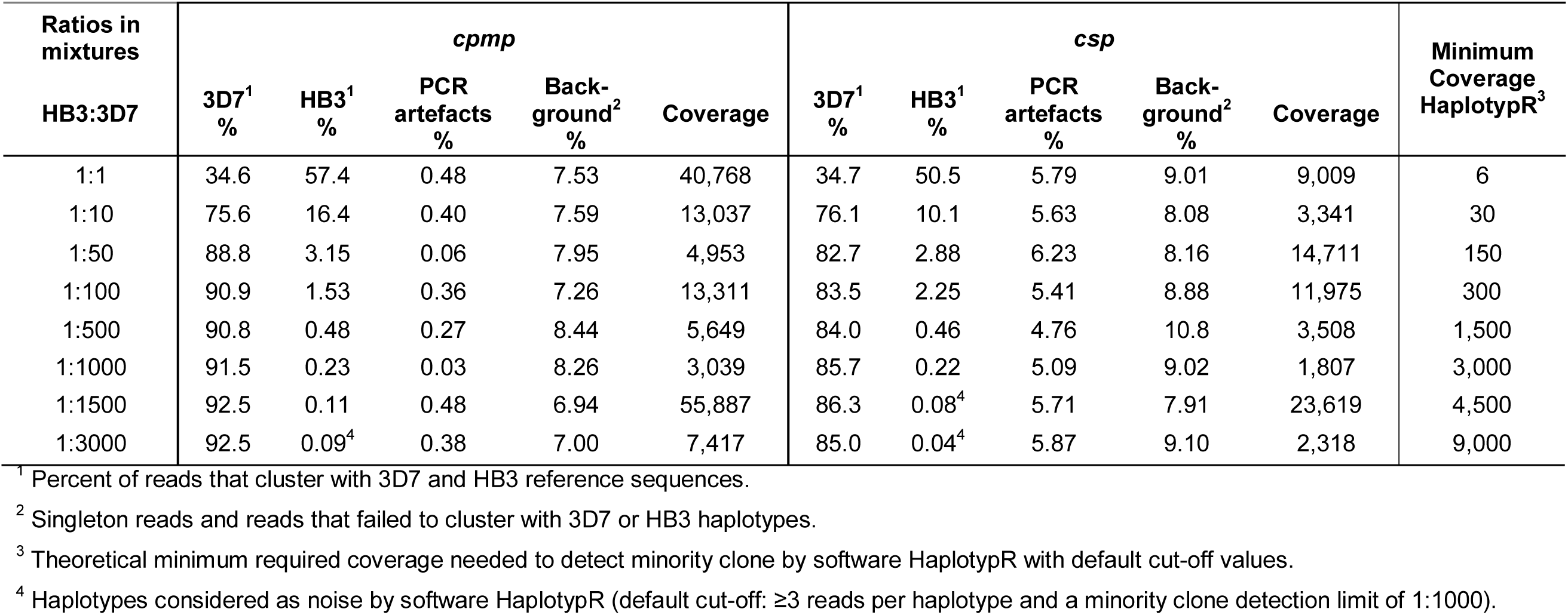
Detectability of the minority clone in defined ratios of *P. falciparum* strains HB3 and 3D7.

| Ratios in mixtures | cpmp |  |  |  |  | csp |  |  |  |  | Minimum Coverage HaplotypR<sup>3</sup> |
| --- | --- | --- | --- | --- | --- | --- | --- | --- | --- | --- | --- |
| HB3:3D7 | 3D7<sup>1</sup> % | HB3<sup>1</sup> % | PCR artefacts % | Back-ground<sup>2</sup> % | Coverage | 3D7<sup>1</sup> % | HB3<sup>1</sup> % | PCR artefacts % | Back-ground<sup>2</sup> % | Coverage |  |
| 1:1 | 34.6 | 57.4 | 0.48 | 7.53 | 40,768 | 34.7 | 50.5 | 5.79 | 9.01 | 9,009 | 6 |
| 1:10 | 75.6 | 16.4 | 0.40 | 7.59 | 13,037 | 76.1 | 10.1 | 5.63 | 8.08 | 3,341 | 30 |
| 1:50 | 88.8 | 3.15 | 0.06 | 7.95 | 4,953 | 82.7 | 2.88 | 6.23 | 8.16 | 14,711 | 150 |
| 1:100 | 90.9 | 1.53 | 0.36 | 7.26 | 13,311 | 83.5 | 2.25 | 5.41 | 8.88 | 11,975 | 300 |
| 1:500 | 90.8 | 0.48 | 0.27 | 8.44 | 5,649 | 84.0 | 0.46 | 4.76 | 10.8 | 3,508 | 1,500 |
| 1:1000 | 91.5 | 0.23 | 0.03 | 8.26 | 3,039 | 85.7 | 0.22 | 5.09 | 9.02 | 1,807 | 3,000 |
| 1:1500 | 92.5 | 0.11 | 0.48 | 6.94 | 55,887 | 86.3 | 0.08<sup>4</sup> | 5.71 | 7.91 | 23,619 | 4,500 |
| 1:3000 | 92.5 | 0.09<sup>4</sup> | 0.38 | 7.00 | 7,417 | 85.0 | 0.04<sup>4</sup> | 5.87 | 9.10 | 2,318 | 9,000 |

<sup>1</sup> Percent of reads that cluster with 3D7 and HB3 reference sequences.593 <sup>2</sup> Singleton reads and reads that failed to cluster with 3D7 or HB3 haplotypes.594 <sup>3</sup> Theoretical minimum required coverage needed to detect minority clone by software HaplotypR with default cut-off values.595 <sup>4</sup> Haplotypes considered as noise by software HaplotypR (default cut-off: ≥3 reads per haplotype and a minority clone detection limit of 1:1000).

Simulations by bootstrap resampling were applied to estimate the average detectability of the minority clone at increasing sequencing coverage and decreasing ratios of the minority clone in the 2-strain mixture. Resampling was repeated 1000 times and included only sequence data from mixed strains with >3000 reads coverage. At a coverage of 10’000 sampled reads the minority clone was detected correctly at ratios 1:1 to 1:1000 for *cpmp* and up to 1:500 for *csp* (Figure 3). The cut-off set for haplotype positivity required that a haplotype was detected ≥3 times and represented ≥0.1% of all reads from the respective blood sample. More stringent criteria to call a haplotype (i.e. a higher minimum number of reads) would require a higher coverage for the detection of minority clones. Thus, more stringency in haplotype definition on the one hand reduces sensitivity, but increases specificity by eliminating false haplotypes attributable to background noise (Figure S6 and S7).

**Figure 3:**
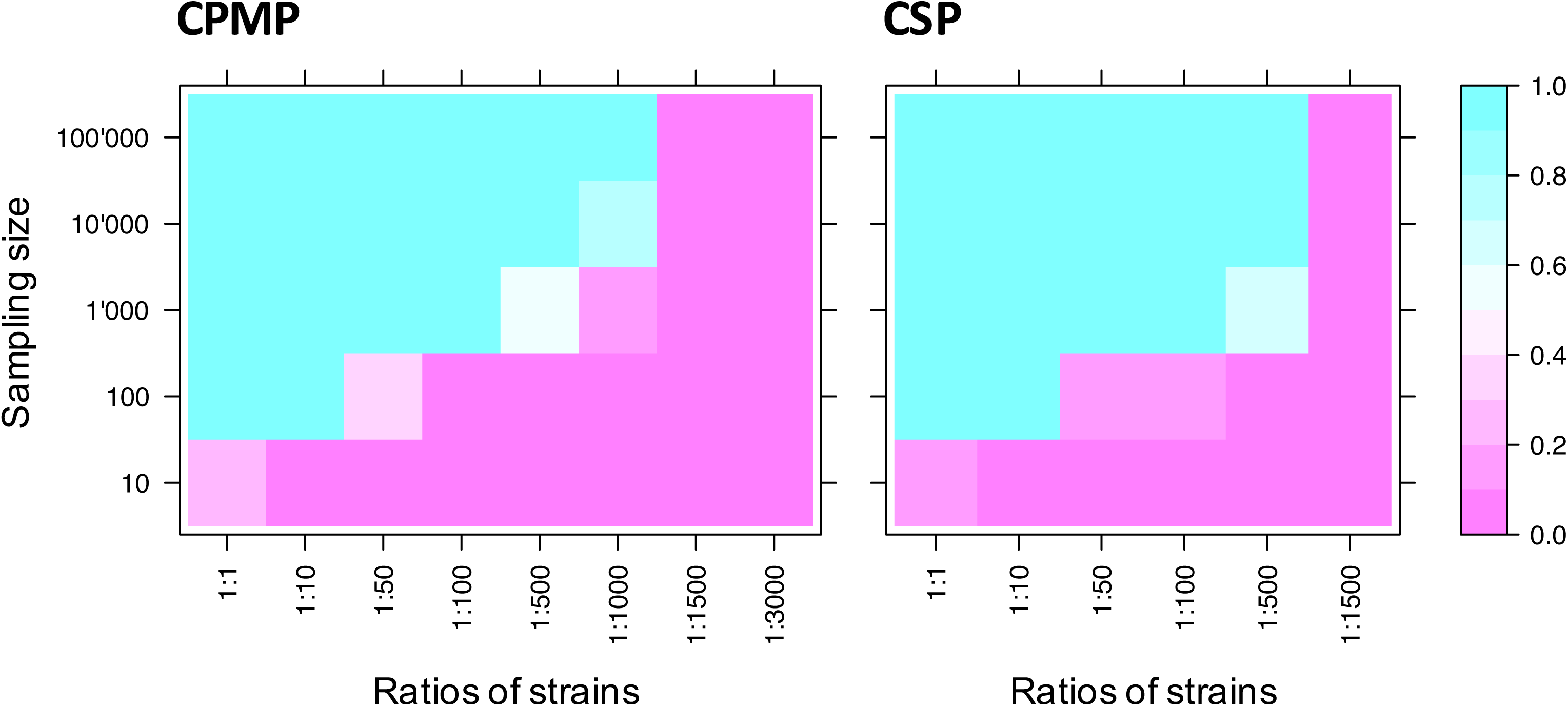
Simulation of minority clone detectability by bootstrapping for marker *cpmp* and *csp*. Cutoff for acceptance of a haplotype was a minimum coverage per haplotype of 3 reads and a minority clone detection limit of 1:1000. Samples were drawn from reads of defined mixtures of *P. falciparum* strain 3D7 and HB3. X-axis shows dilution ratios of strains 3D7 and HB3; Y-axis indicates the sampling size (number of draws from sequence reads) for each mixture of strains. Sampling was repeated 1000 times to estimate mean minority clone detectability.

### Specification of default cut-off settings in software HaplotypR

Cut-off values for the analysis of sequencing data were defined to support removal of background noise caused by sequencing and amplification errors. The following values represent minimal stringency and can be adjusted in the HaplotypR pipeline:

i. Cut-off settings for SNP calling were defined by a population-based approach. A SNP was required to be dominant (>50% of all reads) in ≥2 samples. A single dominant occurrence of a SNP is likely caused by amplification or sequencing error.
ii. Cut-off settings for haplotype calling required a haplotype to be supported by ≥3 reads in ≥2 samples (including independent replicates of the same sample). Per haplotype a minimum of 3 reads are needed to distinguish SNPs from sequencing errors, because a consensus sequence cannot be determined from 2 disparate reads alone. Random sequencing and amplification errors would unlikely lead repeatedly to a particular haplotype.
iii. Cut-off settings for calling minority clones required that a minority clone would represent at least 0.1% of all reads of a sample, which corresponds to a detection limit for minority clones of 1:1000. For the current project, the cut-off was justified by the results obtained from artificial mixtures of culture strains, which defined the technical limit of detection for a minority clone. This parameter may be set to less stringent values.

Application of these three default cut-off values to mixtures of culture strains had the effect that HaplotypR missed the minority clone for both markers in the greatest dilution ration of the two strains tested (1:3000). For marker ***csp*** the minority clone fell below the cut-off even in the 1:1500 ratio (Table 2). No false-positive haplotypes were called.

### Validation of SNP calling

The above criteria were validated on reads from culture strains and primer sequences. The background sequencing error rate at each individual nucleotide position was measured to distinguish sequencing and amplification errors from true SNPs. Mismatch rates of up to 22% was measured in primer sequences (Figure 1), and 18% in amplicons from culture strains (Figure 2, Table S2). None of these mismatches led to calling of a SNP after the above cut-off was applied (i.e. >50% of reads in ≥2 samples).

### Validation of amplicon sequencing in field samples

37 *P. falciparum* samples from PNG were genotyped by amplicon sequencing. Dendrograms were produced for each marker from raw sequencing reads (Figure 4, Figure S8). Branch lengths in these dendrograms represent the number of SNPs that differ between any sequences compared. Branches with sequences belonging to the same haplotype (defined as “clusters”) are labelled in the same colour. Haplotype frequencies within each individual sample were determined from the reads of the sample before applying cut-offs (Figure 4, panel “Quantification”). When analysing the genetic diversity in field sample, haplotypes were only counted as true haplotypes if both replicates pass the haplotype calling cut-off. This more stringent criterion was introduced to prevent erroneous over-estimation of multiplicity due to false haplotypes.

**Figure 4:**
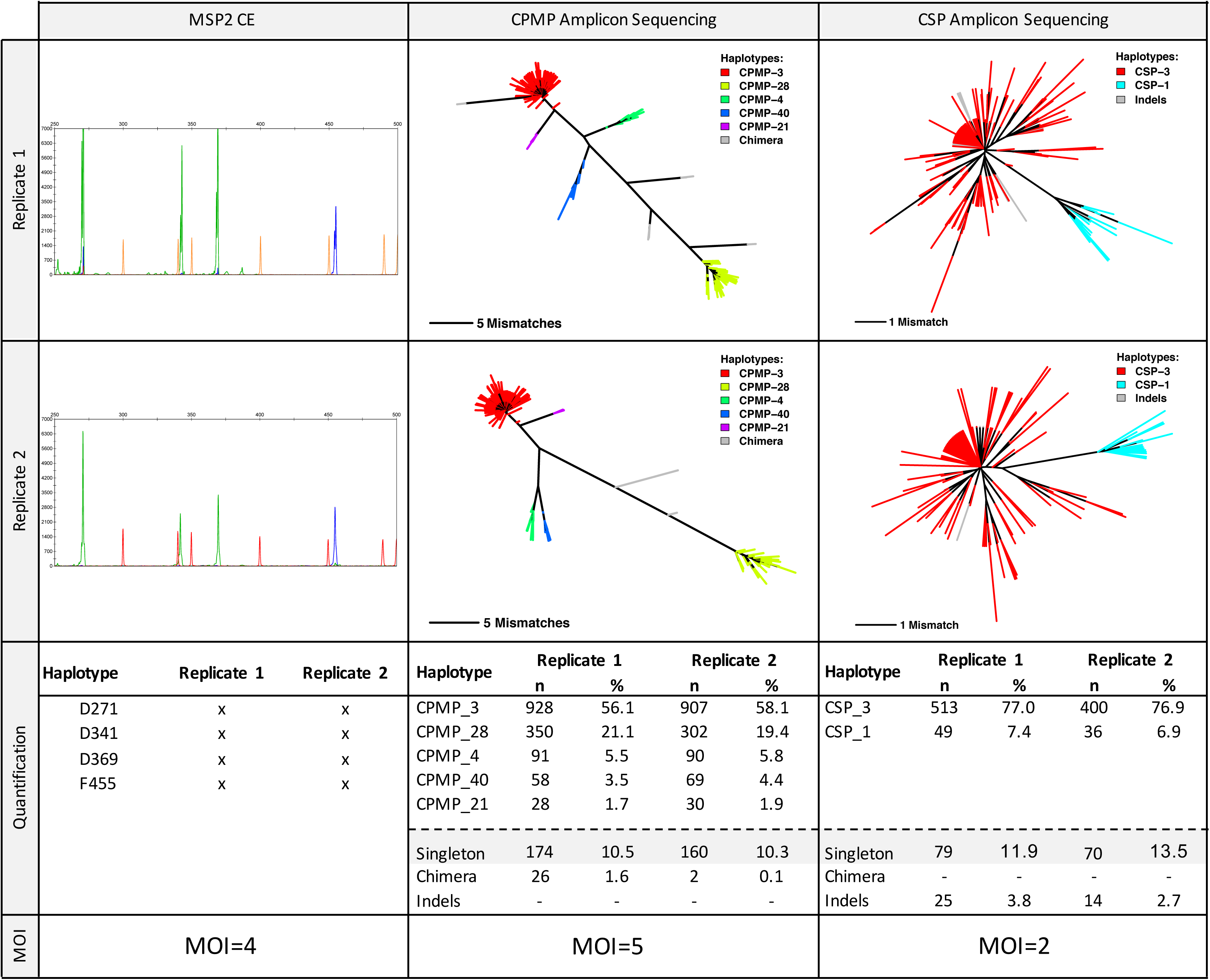
Comparison of genotyping by length-polymorphic marker *msp2* and amplicon sequencing of *cpmp* and *csp*. Raw data for 1 field sample is shown: capillary electropherograms (CE) represent marker *msp2*-CE, dendrograms represent markers *cpmp* and *csp* (two top panels). Quantification of haplotypes and final multiplicity call (two bottom panels). Grey shaded are reads excluded by cut-off settings.

All samples were genotyped for length polymorphic marker *msp2* using capillary electrophoresis (CE) for fragment sizing. *Msp2* genotyping was reproducible and consistent between different laboratories (Figure 4, Figure S8: left column). A mean MOI of 2.2 was observed in 37 field samples analysed by *msp2* genotyping and 25/37 (67.5%) of samples harboured multiple clones (Figure 5C, Table S4). Mean MOI and H_e_ were compared between the genotyping methods (Table 3, Figure 5, Table S4). The resolution of marker *cpmp* was slightly higher than that of *msp2* with 27 *cpmp* haplotypes versus 25 *msp2* alleles, He of 0.96 versus 0.95 and a higher mean MOI of 2.41 versus 2.19, respectively. Overall the two methods agreed well, with good concordance of MOI (Cohen’s Kappa 0.71, equal weights, z=6.64, p-value=3.04e-11). Compared to *msp2* the discriminatory power of *csp* was substantially lower with only 4 *csp* haplotypes found in 37 samples, H_e_ of 0.57 and mean MOI of 1.54. Concordance between *csp* and *msp2* MOI was poor (Cohen’s Kappa 0.38, equal weights, z=4.48, p-value=7.61e-6).

**Figure 5:**
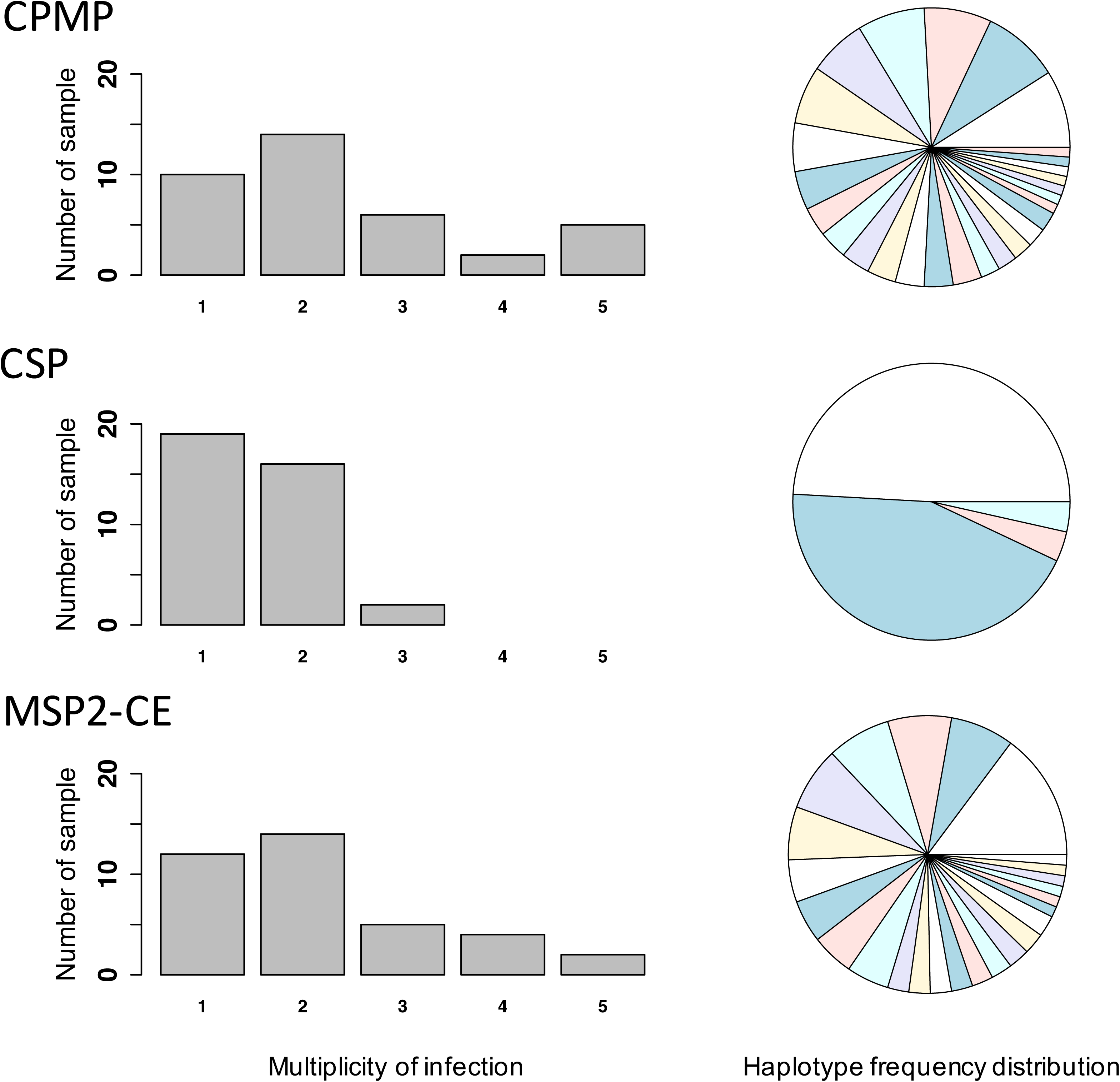
Frequency distribution of multiplicity of infection and allelic frequencies of *cpmp*, *csp* and msp2-CE. 37 field samples were analyzed for each marker.

**Table 3:** Summary of genotyping results from three molecular markers analysed in 37 field samples.

| Marker | $H_e$ | Mean MOI | Number SNPs<sup>1</sup> | Number Haplotypes | Concordance of MOI K |
| --- | --- | --- | --- | --- | --- |
| *msp2* CE | 0.948 | 2.19 | NA | 25 | Reference |
| *cpmp* | 0.957 | 2.41 | 45 | 27 | 0.71<sup>2</sup> (good) |
| *csp* | 0.574 | 1.54 | 10<sup>3</sup> | 4 | 0.38<sup>4</sup> (poor) |

<sup>1</sup> With respect to the reference sequence of *P. falciparum* strain 3D7.
<sup>2</sup> Cohen's Kappa (2 raters, weights=equal): $z = 6.64$ , $p\text{-value} = 3.04e-11$ .
<sup>3</sup> 4/10 SNPs are fixed within these 37 field samples.
<sup>4</sup> Cohen's Kappa (2 raters, weights=equal): $z = 4.48$ , $p\text{-value} = 7.61e-6$ .

### Reproducibility of amplicon sequencing in field samples

*Csp* and *cpmp* haplotypes obtained from 37 field samples were compared between replicates to investigate reproducibility of the molecular and bioinformatic analyses. For both replicates of the field samples the default cut-off criteria for haplotype calling (≥3 reads and minority clone detection limit of 1:1000) were applied. Concordance between replicates was very good with Cohen’s Kappa 0.84 (equal weights, z=7.769, p-value=7.99e-15) for *cpmp* and 0.91 (equal weights, z=6.466, p-value=1.01e-10) for *csp*. Comparison of replicates permitted to investigate the amount of false haplotype calls. True haplotypes should be detected in both replicates, unless the sequence depth is not sufficient for detecting a minority clone in one of the replicates. *Cpmp* minority clones that had a frequency >1.0% of all reads were consistently detected with ≥3 reads in both replicates (Figure S9, Table 4). For *csp* this was achieved for minority clones with a frequency of >0.70%. 18 *cpmp* haplotypes were detected with ≥3 reads in only one of the replicates. In three instances one of the replicate did not pass the cut-off criteria due to low coverage. For marker *csp*, 2 haplotypes with ≥3 reads were detected in one replicate only. In summary, a comparison of replicates indicated 15 potentially false haplotype calls for *cpmp* and 2 for *csp*. These calls stem from reads with a frequency <1%, Therefore, performing replicates are essential to prevent erroneous over-estimation of multiplicity due to false haplotypes.

**Table 4:** Concordance of haplotype calls in replicates of 37 field samples. Bold rows indicate haplotypes that did pass cut-off criteria in both replicates.

|  | *cpmp* | *csp* | Passed cut-off<sup>1</sup> |
| --- | --- | --- | --- |
| Haplotype frequency within sample $\geq 1\%$ | | | |
| **present in both replicates** | **87** | **57** | **yes** |
| present in single replicate only | 0 | 0 | no |
| Haplotype frequency within sample $< 1\%$ | | | |
| **present in both replicates at $\geq 3$ reads<sup>2</sup>** | **2** | **0** | **yes** |
| present in both replicates one $\geq 3$ reads<sup>2</sup> and one $< 3$ reads<sup>2</sup> | 1<sup>3</sup> | 0 | yes/no<sup>4</sup> |
| present in single replicate at $\geq 3$ reads<sup>2</sup> | 17<sup>5</sup> | 2 | yes/no<sup>4</sup> |
| present in both replicates at $< 3$ reads<sup>2</sup> | 1 | 0 | no |
| present in single replicate at $< 3$ reads<sup>2</sup> | 10 | 5 | no |

<sup>1</sup> Default cut-off criteria to accepted haplotype $\geq 3$ reads and a minority clone detection limit of 1:1000.
<sup>2</sup> Owing to default cut-off for haplotype call.
<sup>3</sup> Second replicate had too low coverage to detect $\geq 3$ reads.
<sup>4</sup> Potential false haplotype calls as only one replicate passed cut-off criteria.
<sup>5</sup> In 2 instances second replicate had too low coverage to detect minority clone.

An attempt was made to investigate the influence of the number of PCR cycles performed during amplicon library preparation on the generation of artefacts. This was possible by using 25 and 15 cycles in the nested PCR for replicate 1 and 2, respectively. Cycle number had no influence on the proportion of singleton and indel reads. However, the proportion of chimeric *cpmp* reads was higher in replicate 1 using 25 cycles than in replicate 2 using 15 cycles (0.63% versus 0.13%, Student's t-Test P-value = 0.0221). No chimeric *csp* reads were detected in the field samples (Table 5).

**Table 5:** Mean proportion of singleton or chimeric reads and indels detected in both field sample replicates.

| Marker | Replicate 1 |  |  | Replicate 2 |  |  |
| --- | --- | --- | --- | --- | --- | --- |
|  | Singletons % | Indels % | Chimera % | Singletons % | Indels % | Chimera % |
| *csp* | 11.55 | 3.78<sup>1</sup> | 0.00 | 11.47 | 4.05<sup>1</sup> | 0.00 |
| *cpmp* | 9.76 | 0.073<sup>2</sup> | 0.631<sup>3</sup> | 9.74 | 0.034<sup>2</sup> | 0.130<sup>3</sup> |

<sup>1</sup> Marker *csp*: Indels Replicate 1 versus 2; Student's t-Test: $t = -1.336$ , $df = 71.052$ , $p\text{-value} = 0.1858$
<sup>2</sup> Marker *cpmp*: Indels Replicate 1 versus 2; Student's t-Test: $t = 1.3304$ , $df = 71.94$ , $p\text{-value} = 0.1876$
<sup>3</sup> Marker *cpmp*: Chimera Replicate 1 versus 2; Student's t-Test: $t = 2.3552$ , $df = 55.4$ , $p\text{-value} = 0.02208$

## Discussion

This report presents the development of a new genotyping methodology for *P. falciparum* based on amplicon deep sequencing. The search for new markers was prompted by severe limitations of length polymorphic markers, which represent the currently used standard for genotyping malaria parasites. A strong bias towards preferential amplification of shorter fragments in multi-clone infections was observed, so that larger fragments were lost even if only 5-fold underrepresented compared to shorter fragments from the same sample (7). This called for an alternative approach that relies on haplotypes created from several SNPs rather than length polymorphism. With respect to minority clone detectability, amplicon sequencing overcame this pitfall of length polymorphism methods and also performed very well in field samples.

Amplicon sequencing showed an excellent resolution when using the novel genotyping marker *cpmp* (PF3D7_0104100). The strategy applied for down-selecting highly diverse regions in the genome suggested *cpmp* as the top candidate. *Cpmp* is most abundantly expressed in sporozoite stages (18), but the function of the encoded protein is unknown. The gene is under balancing selection with a Tajima’s D of 1.16 in Guinea and 1.05 in Gambia (19). In this study, *cpmp* revealed a genetic diversity similar to the length polymorphic region of the widely-used marker *msp2*. 45 SNPs were observed in the 37 field samples of this study, leading to the designation of 27 haplotypes for marker *cpmp*. With increasing number of field samples processed, additional rare SNPs and even more haplotypes are likely to be found. The diversity of *cpmp* was high also in the global MalariaGEN dataset (H_e_= 0.93); its resolution as genotyping marker in other geographic regions remains to be shown. In contrast, marker *csp*, analysed in parallel to *cpmp* and also used in earlier studies, showed a limited diversity with only 4 haplotypes detected in 37 field samples. Earlier studies reported similar low diversity for *csp* in regions of Asia Pacific (20). Thus, *csp* is not suited to serve as a single genotyping marker in PNG. However, the global diversity of *csp* according to the MalariaGEN dataset seems to be high (H_e_= 0.86), and high diversity has also been observed in African isolates (20).

Implementing amplicon sequencing required parallel development of a bioinformatics pipeline. A known problem in sequence analysis is the robust detection of minority clones from a background of experimentally induced artefacts. We addressed this problem with the design of HaplotypR, a software package dedicated to stepwise analyse of sequence reads for samples containing multiple clones. The HaplotypR pipeline can be divided into three steps: In the first step, this pipeline de-multiplexes and clusters raw sequence reads to clusters of related sequences, so called “representative haplotypes”. This step employs Swarm2 software, which uses a *de novo* single-linkage-clustering algorithm (21,22). In short, pools of amplicons (identical sequence reads) are expanded by iteratively joining pools of reads that are separated by a defined number of mismatches (e.g. one substitution, insertion or deletion). This strategy permits unbiased clustering of sequence reads without the need to define a list of SNPs. This enables capturing of previously unknown SNPs without any adjustments to the pipeline. In the next step HaplotypR checks all representative haplotypes for presence of PCR artefacts (indels and chimeras), and labels and censors these. In the final step HaplotypR removes background noise by applying defined cut-offs and reports a list of final haplotypes calls.

Validation of HaplotypR was made possible by reads from serial dilutions of *P. falciparum* culture strain 3D7 and from controlled mixtures of strains HB3 and 3D7. On those control samples the impact of amplification and sequencing errors could be assessed. An increased frequency of sequence mismatches relative to the 3D7 reference sequence of up to 22% was observed at a few specific genomic locations including the sequences of amplified primers. To differentiate these sequencing errors from true genotypes of rare minority clones, we defined a SNP calling cut-off where a genotype was required to be dominant (>50% of all reads) in at least 2 samples. This cut-off is critical to distinguish true positive genotypes that are rare in the population from sequencing errors.

To prevent reporting of false haplotypes, HaplotypR pipeline applies two types of cut-offs: firstly, a cut-off for singleton exclusion, whereby a SNP or haplotype needed to be supported by more than one sample. It is unlikely that these cut-offs would remove true haplotypes, except if the sample size was very small. In this case, it is recommended to amplify and sequence samples in duplicate, as in this study. A true haplotype is expected to be present in both replicates and thus will not get excluded. Secondly, a cut-off for haplotype coverage was defined requiring that a haplotype is supported by a user-defined number of sequence reads. This flexible cut-off can be selected for each marker. The coverage cut-off removes false or weakly supported haplotypes and thus improves specificity. On the other hand, the ability to detect minority clones (i.e. sensitivity) will be limited by a cut-off based on coverage. Sequence reads from a minority clone were detected in all ratios up to 1:3000 in the mixtures of strains HB3 and 3D7. However, due to high background noise, false haplotypes with a frequency of up to 0.01% were also detected, making the definition of a cut-off to remove background noise obligatory. Applying these default cut-offs in HaplotypR decreased minority clone detectability from 1:3000 to 1:1000.

The potential of amplicon sequencing for genotyping samples of very low parasitaemia was assessed in serial dilutions of strain 3D7. Sequence reads were retrieved from samples of a parasitaemia as low as 5 parasites/μl, however coverage was below 100 reads for the lowest level of parasitaemia. To reliably genotype samples spanning a wide range of parasitaemias, similar sequence coverage (and thus unbiased normalization of input material) for all samples is needed. The inexpensive strategy used to adjust amplicon concentrations of individual samples to equal levels prior to pooling for highly multiplexed sequencing still resulted in fluctuation in the sequence coverage, but a commercial DNA normalisation kit may improve equimolar pooling of samples (23,24).

All samples in this study were sequenced in 2 replicates. This was done to assess the reproducibility of amplicon sequencing method of genotyping very low abundant minority clones, and to investigate the effect of nested PCR cycle number on artefacts. Analysing replicates of field samples revealed that haplotypes with a frequency of >1% were consistently detected in both replicates. In contrast, haplotypes with a frequency of <1% were frequently detected only in a single replicate. If minority clones of <1% frequency are to be reliably detected, amplifying and sequencing two or more replicates for each sample would be essential to call true haplotypes.

To detect minority clones with high sensitivity, samples need to be sequenced at high coverage and preferable in replicates. As sensitivity may be adjusted by sequence coverage, choices have to be made in a trade-off between sequencing costs and sensitivity. The specific genotyping application can guide this choice. For example, in large scale field studies with many samples, a high degree of multiplexing of samples at moderate sequence coverage may be chosen to keep sequencing cost low. Furthermore, a less sensitive approach without performing replicates may be sufficient when detection of very rare minority clones is less of an issue. Another important application of genotyping of malaria parasites is the example of “recrudescence typing” during *in vivo* drug efficacy trials. To distinguish a new infection from one present as a minority clone prior to drug administration requires highest sensitivity and every clone must be reliably detected. In such cases a sequencing approach with less multiplexing is desired to achieve high coverage and maximal detection of minority clones.

The power of high sequencing coverage was shown for example in a study assessing the subclonal diversity in carcinomas (25). Minority variants with a frequency of 1:10’000 were detected with a sequence depth of 100’000 reads per sample. Our results reported from malaria field samples does not have sufficient sequence depth to achieve such sensitivity, as median sequencing depth per sample was 1’490 reads for *cpmp* and 731 reads for *csp* owing to a high number of samples and of markers sequenced in parallel. As 384 samples were analysed in a single sequence run following a PCR based sequencing library preparation, costs were comparable to those of previous standard *msp2-CE* genotyping (5). Thus, the approach applied by us is cost effective as it permits parallel processing of several hundred samples, a range typically encountered in population-wide studies.

## Conclusions

Short amplicon sequencing has the advantage that no multi-locus haplotype reconstruction is needed, as all SNPs are linked by a single paired-end read. This allows the reliable analysis of samples of very high MOI, a prerequisite for genotyping in areas of high malaria endemicity. An additional strength of this method is that previously undescribed or newly evolving haplotypes can be captured without any adjustment of the typing methodology or the HaplotypeR pipeline. The main limiting factor for the detection of minority clones was the sequence depth per sample. If detection of clones in a 1’000-fold under-representation compared to a dominant clone is required, studies should aim for a minimum sequence coverage of 10’000 reads per sample and perform experiments in duplicates.

The specification of amplification and sequencing errors presented here as well as the developed bioinformatic tools to handle such complex analytical tasks are relevant to all amplification-based genotyping methods of multiple clones or quasi-species within a sample. The newly developed pipeline can be used to analyse any amplicon sequencing based genotyping data irrespective of marker or organism.

## Methods

### Parasite genomic DNA

*P. falciparum in vitro* culture strains HB3 and 3D7 were mixed in 8 different proportions to generate well defined control samples with known MOI and well defined ratios of genomes. The ratios in these HB3-3D7 mixtures ranged from 1:1 to 1:3’000. Five additional control samples represented a dilution series of strain 3D7 with parasite densities ranging from 50’000 to 5 parasite/μl. Dilutions were prepared in human gDNA to reconstitute the nucleic acid concentration of a human blood sample. Details of parasite quantification were published previously (7). 37 archived field samples from a cohort study conducted in East Sepik Province, Papua New Guinea (PNG) in 2008 were used to validate the performance of protocols for genotyping and data analysis in natural *P. falciparum* infections (26). Ethical clearance was obtained from PNG Institute of Medical Research Institutional Review Board (IRB 07.20) and PNG Medical Advisory Committee (07.34). Informed written consent was obtained from all parents or guardians prior to recruitment of each child.

### Genotyping using length polymorphic marker *msp2*

For determination of mean MOI field samples were genotyped using the classical *P. falciparum* genotyping marker *msp2*. Fluorescently labelled nested PCR products were sized by CE on an automated sequencer and analysed using GeneMapper software according to previously published protocols (5). Each DNA sample was genotyped twice in independent laboratories to assess reproducibility of clone multiplicity (Figure 4 and S6).

### Amplicon deep sequencing marker selection and assay development

3’411 genomes from 23 countries, published by the *Plasmodium falciparum* Community Project (MalariaGEN), were screened to identify highly diverse markers for SNP-based genotyping (16). The *P. falciparum* genomes were divided in 200bp windows and H_e_ was calculated for each window as follows: 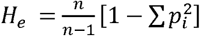 where *n* is the number of clones and *p_i_* the frequency of allele *i*. Annotated genes (PlasmoDB v11.0) that overlapped with windows of high heterozygosity were selected for further evaluation. Genes belonging to gene families, such as *var, rifin, stevor* and surf families, were excluded from the list, as well as genes with high heterozygosity that is usually caused by length polymorphism (Supplementary File S2).

Primers for marker *cpmp* were designed manually. Location of primers was selected to flank a region of maximum diversity (Figure S1, Figure S2). Amplicon sizes were limited to a maximum of 500bp to conform to possible read lengths of the Illumina MiSeq platform. Quality control of primers was assessed with online tools for secondary structure and primer dimer interaction (27,28). Primer sequences are listed in Table S1 and H_e_ values for amplicons are shown in Supplementary Figures S1 and S2.

### Sequencing library preparation

The sequencing library was generated by 3 rounds of PCR with KAPA HiFi HotStart ReadyMix PCR Kit. A first round of 25 cycles amplified the gene of interest. A second marker-specific nested PCR amplified the primary product with primers that carried a 5’ linker sequence. We compared different cycle numbers for this second round: 25 cycles for replicate 1 and 15 cycles for replicate 2. This comparison was done to test for effects of cycle number on sequence diversity caused by imperfect polymerase fidelity (29). To allow pooling and later de-multiplexing of amplicons, a third and final amplification was performed using primers binding to the F and R Linker sequence at the 3’ end, that introduced a sample-specific molecular barcode sequence plus the Illumina sequence adapter at the 5’ end. The relative positions of all these elements are depicted in the schematic in Figure S4. A detailed PCR protocol containing primer sequences, cycle conditions und pooling steps are described in Supplementary File S1 and Table S1.

PCR products were purified with NucleoMag beads. The expected fragment size of the sequencing library was confirmed by Agilent 2200 Tapestation System. DNA concentration of the sequencing library was quantified by Qubit Fluorometer (Thermo Fisher Scientific). Sequencing was performed on an Illumina MiSeq platform in paired-end mode using Illumina MiSeq reagent kit v2 (500-cycles) together with a Enterobacteria phage PhiX control (Illumina, PhiXControl v3).

### Bioinformatic analysis pipeline “HaplotypR”

Sequence reads were mapped with bowtie2 (30) to the *phiX174* genome (Accession: J02482.1) for assessing the quality of the sequencing run and calculating sequencing error per nucleotide position. Reads were then de-multiplexed to separate individual samples and different genotyping markers (Figure 6). Primer sequences were truncated, the sequence was trimmed according to the quality of the *phiX* control sequence reads and paired reads were fused together.

**Figure 6:**
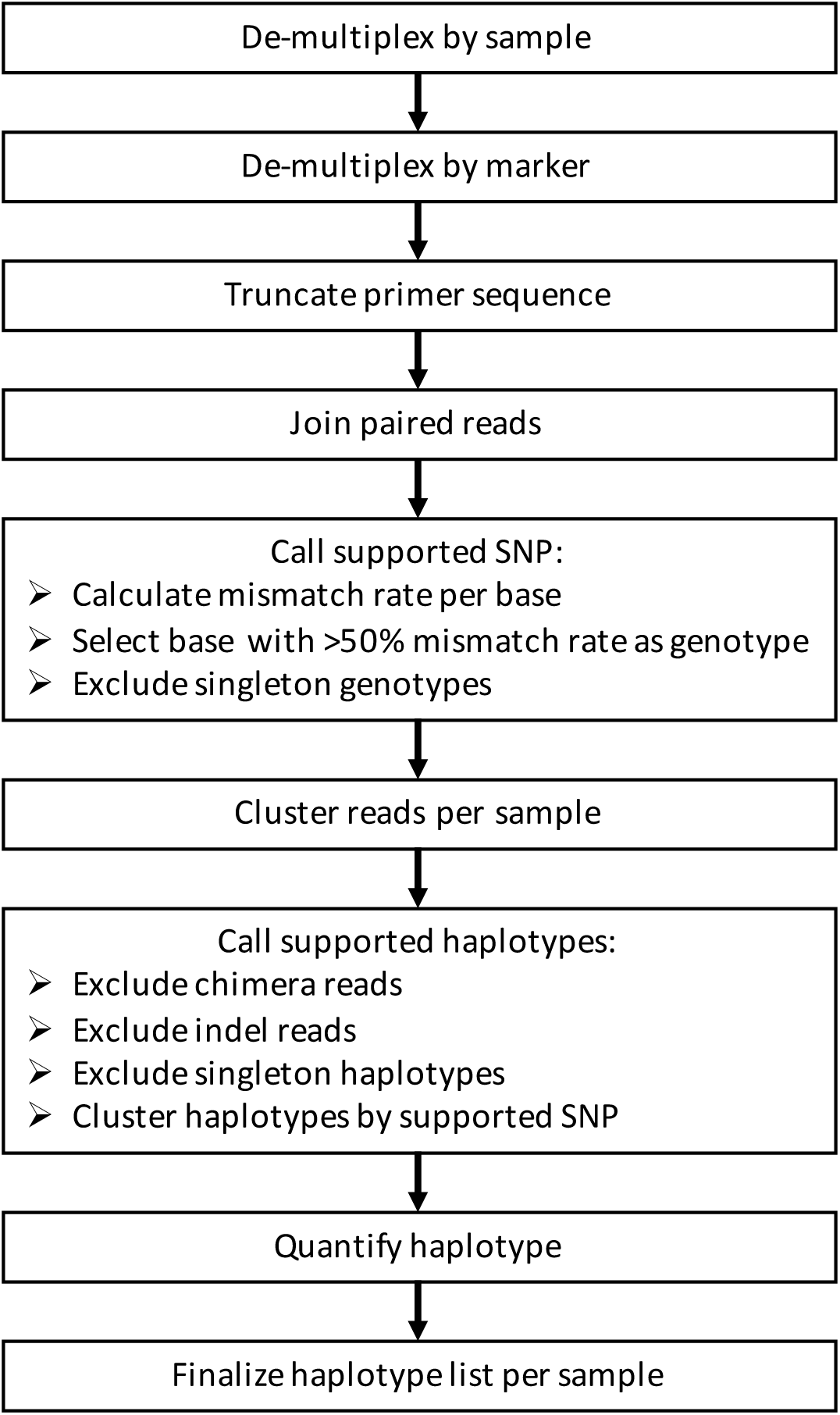
Bioinformatic analysis pipeline applied on highly multiplexed deep sequencing data.

For analysis of control samples, fused reads were mapped to the corresponding primers and *P. falciparum* reference sequences of strains 3D7 and HB3 (Accession: AL844502.2, AL844501.2, AB121018.1, AANS01000117.1). Rates of mismatches to primer and reference sequences were calculated for each individual sample at each nucleotide position. A SNP was defined as a nucleotide position with a >50% mismatch rate in the sequence reads from at least two independent samples.

For prediction of haplotypes, fused reads were clustered individually per sample with Swarm2 software (parameters: boundary=3 and fastidious mode) (21,22). The centre of each cluster represents the most abundant sequence of the cluster and thus constitutes a predicted haplotype. The cluster size represents the within-sample clone frequency in the tested sample. Haplotypes were checked for PCR artefacts such as indels and chimeric reads. Indels are caused by polymerase slippage which occurred primarily at stretches of homo-polymers. Chimeric reads, caused by incomplete primer extension and inhomologous re-annealing, were identified with vsearch software (parameters: uchime_denovo mode, mindiffs=3, minh=0.2) (31). Haplotypes with a cluster size of 1 were classified as singletons and considered background noise. The full analysis pipeline, named HaplotypR, was implemented as R package and is illustrated in Figure 6 (https://github.com/lerch-a/HaplotypR.git).

### Estimated detectability of minority clones by sampling

Detectability of minority clones was estimated by bootstrapping from the reads of the control samples with defined HB3-3D7 strain ratios. Reads were randomly sampled with replacement until the required coverage was reached. These resampled set of reads were processed in the same manner as the original samples using HaplotypR. For resampling only sequence files from HB3-3D7 mixtures were used that had a coverage of >3000 reads.

## Supporting Information

File S1. Supporting information. Supplementary text, figures and tables. (PDF)

File S2. List of 200bp H_e_ windows of whole *P. falciparum* genome. (XLSX)

## Acknowledgement

We are grateful to the study participants and their guardians and to the field and laboratory team of the PNG Institute of Medical Research, in particular Anna Rosanas-Urgell and Alice Ura.

## Authors Contributions

Conceived and designed the experiments: IF, IM, AL, CK, SW

Performed the experiments: AL, CK, JHK, NH, CM, SW

Provided samples: IB

Analyzed the data: AL

Supervision: IF, IM, LOC

Writing - draft: AL, IF

Writing - review & editing: CK, NH, JHK, IM, LOC

